# Extensive novel diversity and phenotypic associations in the dromedary camel microbiome are revealed through deep metagenomics and machine learning

**DOI:** 10.1101/2024.02.05.578850

**Authors:** Fathi A Mubaraki

## Abstract

The dromedary camel, also known as one-humped camel or Arabian camel, is an iconic and economically important feature of Arabian society. Its contemporary importance in commerce and transportation, along with the historical and modern use of its milk and meat products for dietary health and wellness, make it an ideal subject of scientific scrutiny. The gut microbiome has recently been associated with numerous aspects of health, diet, lifestyle, and disease in livestock and humans alike, as well as serving as an exploratory and diagnostic marker of many physical characteristics. Our initial pilot analysis of 55 camel gut microbiomes from the Fathi Camel Microbiome Project using deep metagenomic shotgun sequencing reveals substantial novel species-level microbial diversity, for which we have generated an extensive catalog of prokaryotic metagenome-assembled microorganisms (MAGs) as a foundational microbial reference database for future comparative analysis. Exploratory correlation analysis exhibits substantial correlation structure among the collected subject-level metadata, including physical characteristics. Machine learning using these novel microbial markers, as well as statistical testing, demonstrates strong predictive performance of microbes to distinguish between multiple dietary and lifestyle characteristics in dromedary camels. We derive strongly predictive models for camel age, diet (including wheat), level of captivity. These findings and resources represent substantial strides toward understanding the camel microbiome and pave the way for a deeper understanding of the camel microbiome, as well as the nuanced factors that shape camel health.

## Introduction

The study of the gut microbiome has emerged as a pivotal field of research with far-reaching implications for health and disease across a broad spectrum of hosts, from humans to agriculturally significant mammals. Recent advances have illuminated the deep connections between the microbiome and numerous health outcomes in both human and animal populations, positioning the microbiome as a potential biomarker for various physical traits [1,2,3]. The Human Microbiome Project (HMP) Consortium has notably contributed to this understanding, unveiling the dramatic differences in microbial populations across diverse bodily habitats, even among healthy individuals [4]. Their research underscores the distinction between alpha diversity, the variety of microbes within individual samples, and beta diversity, the differences in microbial communities between different samples [5]. The gut microbiome’s profound implications for health, both in humans and animals, are underscored by the growing body of research that connects its composition and functionality to a host’s wellbeing. Distinctions in alpha diversity—the variety within a particular microbiome sample—and beta diversity—the differences between different microbiome samples— suggest that the microbiome’s effect on health is not uniform but rather specific to each individual’s internal and external ecological environments. This nuanced understanding has implications for personalized medicine and animal husbandry, highlighting the need for tailored approaches in managing health and disease.

Furthermore, the microbiome’s role in animal husbandry has attracted attention, particularly in the context of dietary influences on microbial communities. A study examining the effect of diet on dairy cows highlighted a significant shift in the rumen’s microbial ecosystem when transitioning between pasture grazing and a total mixed ration diet [6]. This work underlines the profound impact that dietary practices have on the microbial consortia within the gastrointestinal tract. Diet’s impact on the microbiome is exemplified in studies of dairy cows, where shifts in the microbial community are evident depending on whether the animals are pasture-fed or given a total mixed ration [7]. These shifts are not merely of academic interest; they have practical implications for agricultural practices and animal nutrition. Understanding the microbiome’s response to different dietary regimens can lead to optimized feeding strategies that improve the health and productivity of livestock.

The Bactrian camel, Camelus bactrianus, with its distinctive pair of dorsal humps, has also been the subject of microbiome research. Investigations into the digestive systems of Bactrian camels using 16S rRNA gene amplicon sequencing have revealed differences in microbial communities; however, these studies have their limitations, as they focus on a single hypervariable region, V4, potentially providing an incomplete picture of the gastrointestinal microbiota [8].

The gut microbiome’s functional contributions are diverse, encompassing roles in food breakdown and energy production for the host. For instance, a focused research effort revealed variabilities in the fecal microbiome of Arabian camels, pinpointing gender as a factor associated with differences in both the composition and functional capacity of the stool microbiome [9].

While machine learning has revolutionized data analysis across various fields, its application to microbiome research has been largely concentrated on human subjects. This leaves a significant gap in the understanding of animal microbiomes. One study demonstrated the utility of machine learning by training several models to predict colonic diseases using human fecal 16S rRNA data [10]. Another employed metagenomic sequencing to probe the relationship between the gut genes of premature infants and their survival strategies in response to specific clinical and environmental conditions, revealing that formula feeding correlates with an increased presence of certain antibiotic resistance genes in the infant gut microbiome [11]. Machine learning presents a powerful tool to decipher the vast and complex datasets generated by microbiome research. While its application has been largely concentrated on human subjects [12], the methodology holds equal promise for animal microbiome studies. In humans, machine learning models trained on fecal 16S rRNA data have shown potential in predicting diseases such as colorectal cancer. Similarly, the use of these models to analyze metagenomic data in infants has shed light on how diet, specifically formula feeding, can impact the gut microbiome’s resistance to antibiotics. These examples underscore the potential of machine learning to reveal patterns that might not be discernible through traditional statistical analyses and could be pivotal for advancing our understanding of the microbiome’s role in animal health and disease.

Despite all this, the Arabian camel, also known as the dromedary camel, remains relatively understudied, especially given its historical and ongoing significance in Arabian societies for transportation and as a source of milk and meat. Notably, the annual camel festival in Saudi Arabia, which celebrates this iconic animal, provides a unique opportunity for scientific observation and research, contributing to the cultural and scientific narrative of these important creatures.

This project seeks to delve into the microbial composition of Arabian camels, advancing our understanding of the diversity within their microbial communities. By leveraging advanced metagenomic shotgun sequencing, data mining, statistical analysis, and machine learning approaches, we aim to explore the camel microbiome comprehensively. This integrative approach will pave the way toward a nuanced appreciation of the factors influencing camel health and, by extension, the welfare of the human communities that depend on them.

## Materials and Methods

### Camel fecal specimen collection and handling

55 camels were sampled from the Tabuk region of Saudi Arabia from a variety of lifestyles and regional topographies. Sampling was performed under supervision 27 distinct herds of camels were selected for our investigation, including up to 4 camels per herd (avg 2.04 / herd). In brief, the sampling protocol consisted of waiting until camel(s) within a herd dropped feces, then immediately collecting top-most scrapings of the final pellet of feces from the excrement and suspending 0.5-0.7g of material with forceps into 99% ethanol buffer within a sealed pre-labeled specimen tube.

### Sample storage and processing

Samples were stored under 40C refrigeration while collection took place over a period of 30 days. Samples were shipped to BGI Hong Kong for DNA extraction and sequencing using the standard BGI complete DNA microbiome extraction kit. The extracted DNA was then sequenced using the DNBseq platform at 2×150bp at 55M sequencing pair read depth (110M total reads per sample). Data was transferred via AWS S3 from BGI to our server for downstream analysis.

### Raw data analysis

QC was performed on the raw sequencing data using SHI7 using default settings [13]. Both a single-sample and pooled assembly was performed; in order to combine data from both methods, species-level representative genomes from the pooled assembly were only added where a single-sample-assembled genome of the same species was not assembled. Assembly was performed using megahit v1.2.9 [14]. MAGs were identified and binned using metabat2 (single sample assembly and, separately, all 55 samples co-assembled on a high-RAM node), and quality was assessed using CheckM2 v1.0.2 [15]. MAGs with assessed completeness > 50% and contamination <= 5% were retained and clustered at 95% ANI in R from a distance matrix formed using aKronyMer v1.0 [16] using ANI GC LOCAL distance parameters with K = 13. Representatives were selected for each cluster with the highest aggregate completeness and lowest contamination (score = completeness - (5*contamination)).

Taxonomy was assigned with GTDB-tk v2.3.0 with GTDB R214 [17]. The resulting set of MAGs were used to create an XTree database for downstream analysis and profiling [18]. Read counts and unique coverage profiles were generated by XTree for all 55 samples, and the resulting representative species-level profiles were compiled into a species-level taxonomy table. To assign a unique species-level placeholder name when GTDB was unable to assign a reference species name, we used the genome ID as a placeholder. Genus-level aggregation was also performed to assign a consistent placeholder name to species-level representatives from species that clustered together at >= 90% ANI (but < 95%) using the same approach outlined above.

### Statistical and machine learning analysis

Correlations were performed in R (v4.3.0). Alpha diversity was computed by summing the number of non-zero species relative abundances per sample (species richness), where a species whose genome is greater than 25% uniquely covered is considered present. Beta diversity was calculated using cmdscale on log10-scaled species relative abundances (log euclidean). Machine learning was performed with the randomForest package in R using the relative abundance data against parameters from the metadata, including dietary and lifestyle features. Reported ML performance scores (AUC for binary features) were calculated using random forest out-of-bag predictions; feature importance was assessed using the Gini index. For increased generalizability in this 55-sample dataset, genus-level taxonomy was used for all machine learning and differential analysis.

## Results

The Fathi Camel Microbiome Project resulted in the collection and metagenomic sequencing of fecal material from 55 camels, along with a number of metadata fields (covariates) per camel. This allowed for the generation of a robust database of metagenome-assembled genomes (MAGs), as well as the ability to perform statistical associations.

### Demographic summary highlights the diversity of sampling

To reflect potential metagenomic diversity in camel microbiomes, we sampled from a diverse set of camels from the Tabuk region of Saudi Arabia. The demographic summary of animal characteristics is presented in Table 1 below, highlighting a diversity of ages, diets, and habitats across 27 herds.

**Table 1.**
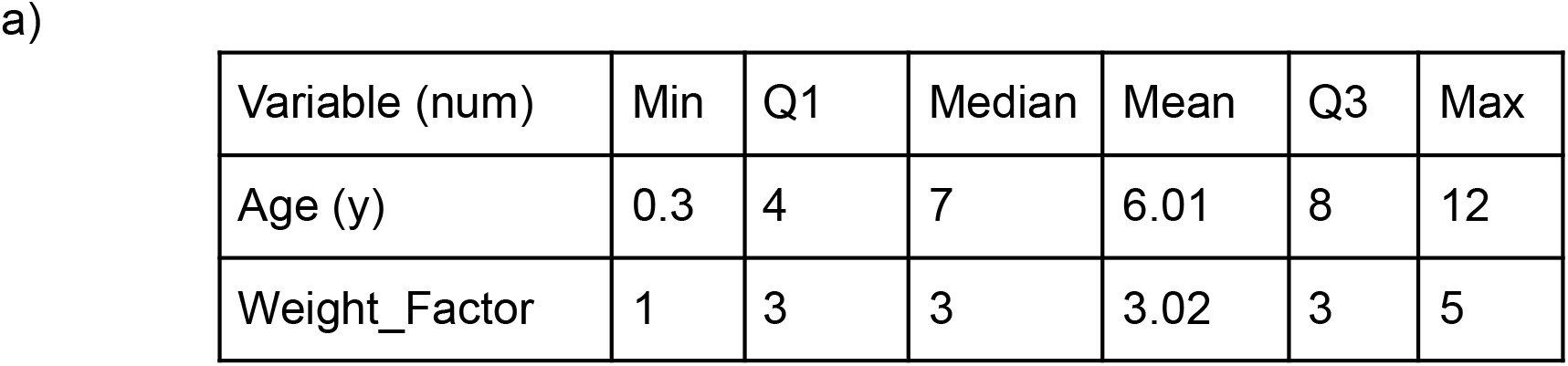
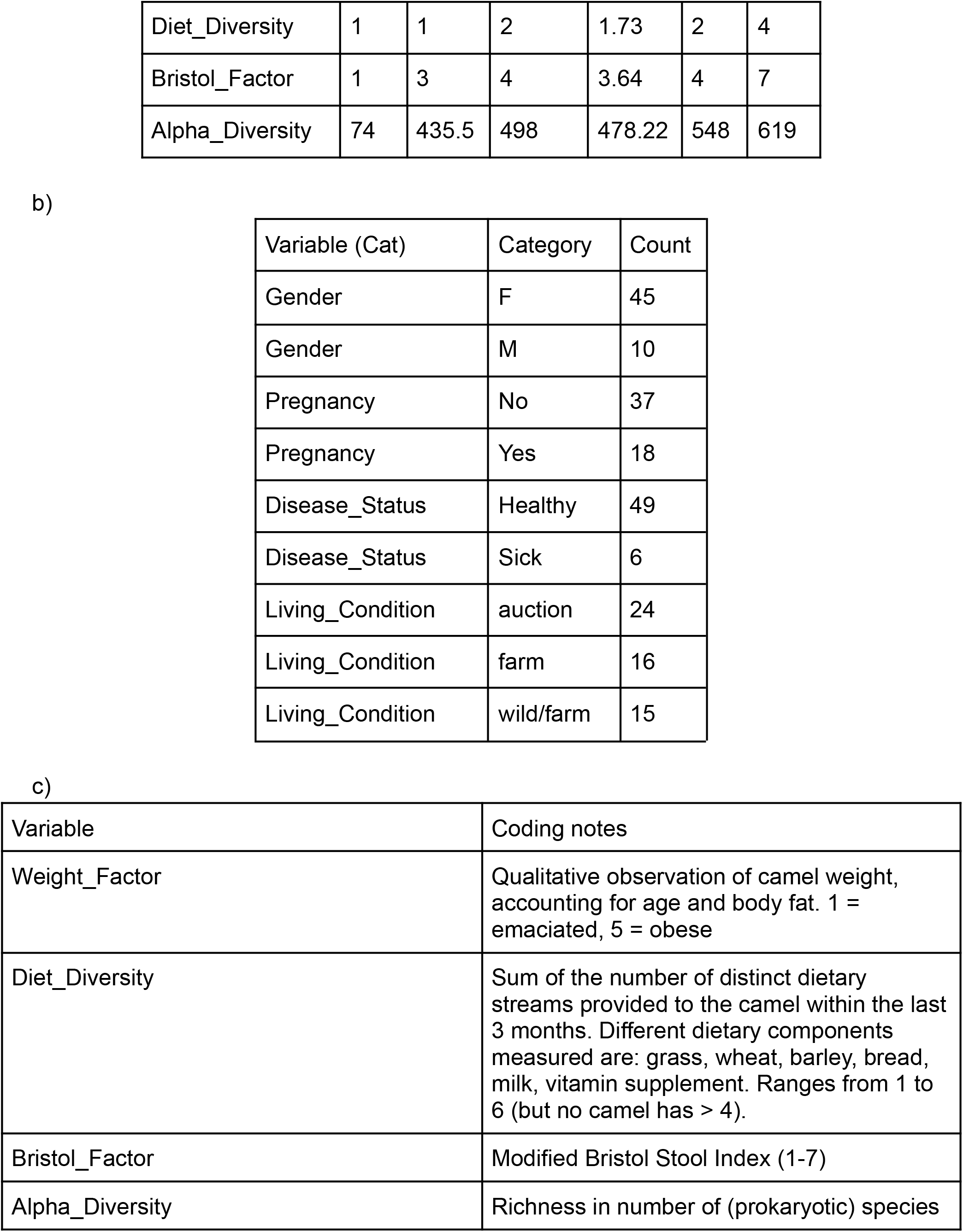
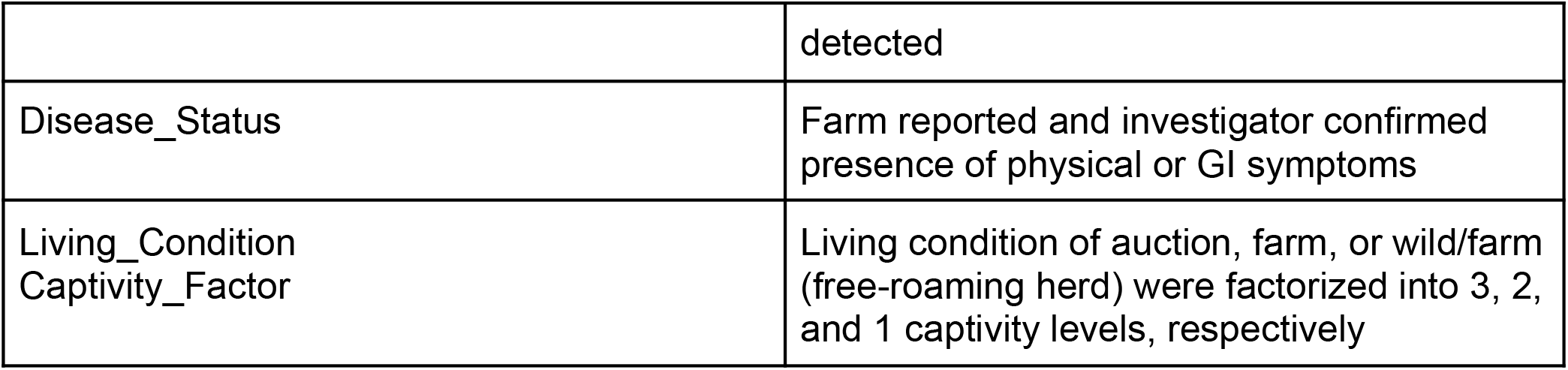
Demographic characteristics of the FCMP, separated into numerical variables (a) and categorical variables (b). Coding was determined as specified in (c). More collected variables are provided in the supplementary metadata table.

| Variable (num) | Min | Q1 | Median | Mean | Q3 | Max |
| --- | --- | --- | --- | --- | --- | --- |
| Age (y) | 0.3 | 4 | 7 | 6.01 | 8 | 12 |
| Weight_Factor | 1 | 3 | 3 | 3.02 | 3 | 5 |
| Diet_Diversity | 1 | 1 | 2 | 1.73 | 2 | 4 |
| Bristol_Factor | 1 | 3 | 4 | 3.64 | 4 | 7 |
| Alpha_Diversity | 74 | 435.5 | 498 | 478.22 | 548 | 619 |

b)
| Variable (Cat) | Category | Count |
| --- | --- | --- |
| Gender | F | 45 |
| Gender | M | 10 |
| Pregnancy | No | 37 |
| Pregnancy | Yes | 18 |
| Disease_Status | Healthy | 49 |
| Disease_Status | Sick | 6 |
| Living_Condition | auction | 24 |
| Living_Condition | farm | 16 |
| Living_Condition | wild/farm | 15 |

| Variable | Coding notes |
| --- | --- |
| Weight_Factor | Qualitative observation of camel weight, accounting for age and body fat. 1 = emaciated, 5 = obese |
| Diet_Diversity | Sum of the number of distinct dietary streams provided to the camel within the last 3 months. Different dietary components measured are: grass, wheat, barley, bread, milk, vitamin supplement. Ranges from 1 to 6 (but no camel has > 4). |
| Bristol_Factor | Modified Bristol Stool Index (1-7) |
| Alpha_Diversity | Richness in number of (prokaryotic) species |
|  | detected |
| Disease_Status | Farm reported and investigator confirmed presence of physical or GI symptoms |
| Living_Condition<br>Captivity_Factor | Living condition of auction, farm, or wild/farm (free-roaming herd) were factorized into 3, 2, and 1 captivity levels, respectively |

### A comprehensive prokaryotic genome database for camel microbiome analysis

An important contribution of this work is the generation of a prokaryotic reference genome database from direct assembly of deeply sequenced camel fecal metagenomes. A total of 3,165 species-level prokaryotic (bacterial and archaeal) genomes were produced with CheckM2 completness >= 50% and contamination <= 5%, per MiMAG quality criteria. After taxonomy assignment, 726 genera were found, with 55 containing > 10 species. Up to 151 genera were novel (as they contained no CheckM2 identification). 2,740 of the 3,165 species-level genomes (87%) represented novel species without any species-level database reference or representative genome in the GTDB. This high level of novelty is expected given the lack of deep metagenomic sequencing efforts in dromedary (Arabian) camels until the current study.

### Associations between metadata and diversity

A pairwise all-vs-all spearman correlation among continuous variables showed some expected correlations between collection-related variables, such as time of day and temperature (Figure 2a). Some biological variables also clustered by correlation, such as Bristol index (stool consistency) with stool darkness in one cluster, and alpha diversity with age in another. As expected, level of captivity inversely correlated with the amount of grazing reported, as well as alpha diversity and age (younger camels are more likely to be kept in controlled auction environments).

**Figure 1.**
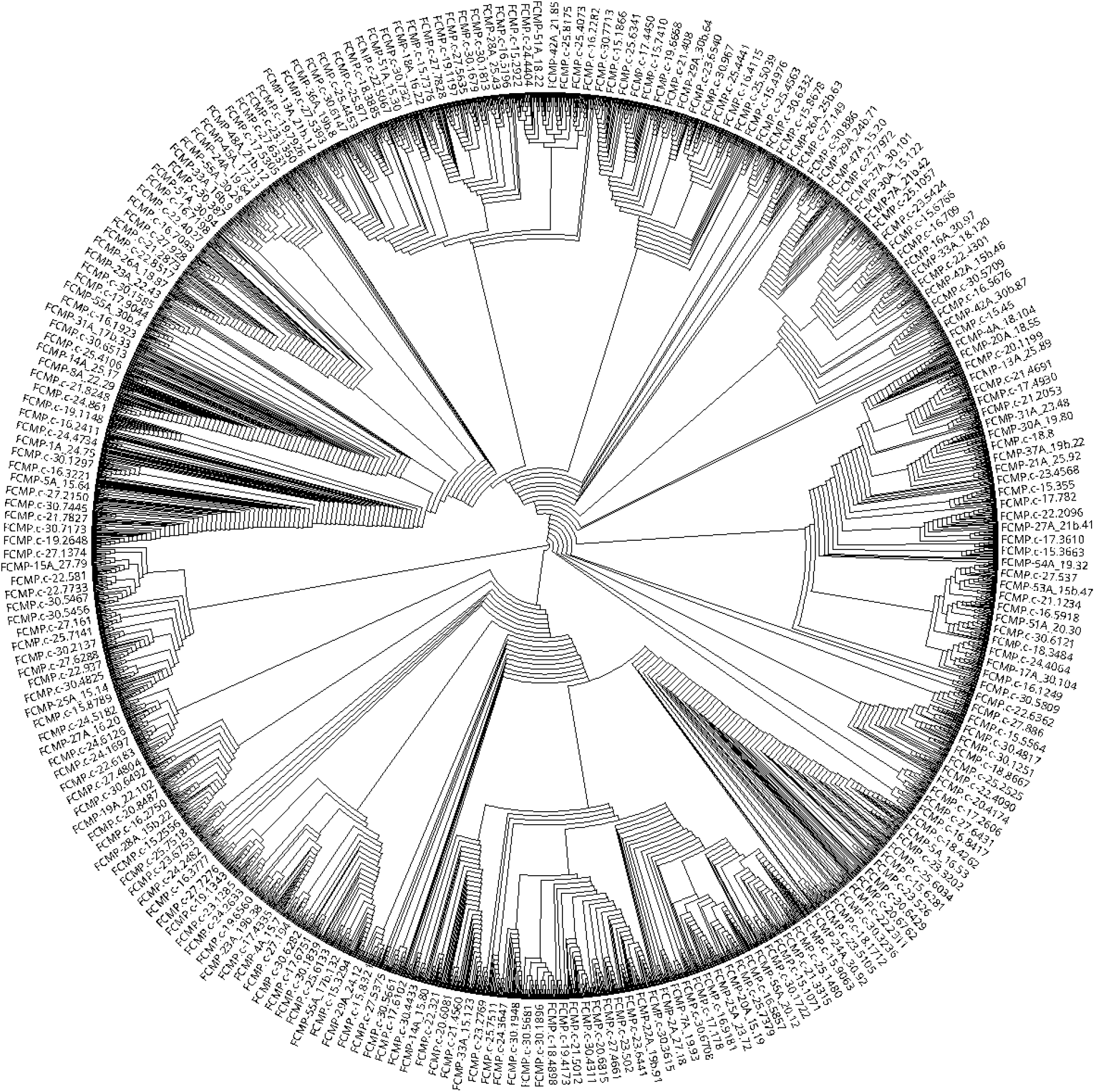
An extensive set of reference genomic MAGs spanning 3,165 new species-level representatives. Average nucleotide identity-based ladderized circular cladogram depicting approximate relationship among the genomes. Narrow, deeply-branched regions indicate phylogenetic singleton genomes, while wider clades near the edges indicate the presence of more closely-related species. Sparse labels on the exterior comprise an even sampling of diversity along the tree.

**Figure 2.**
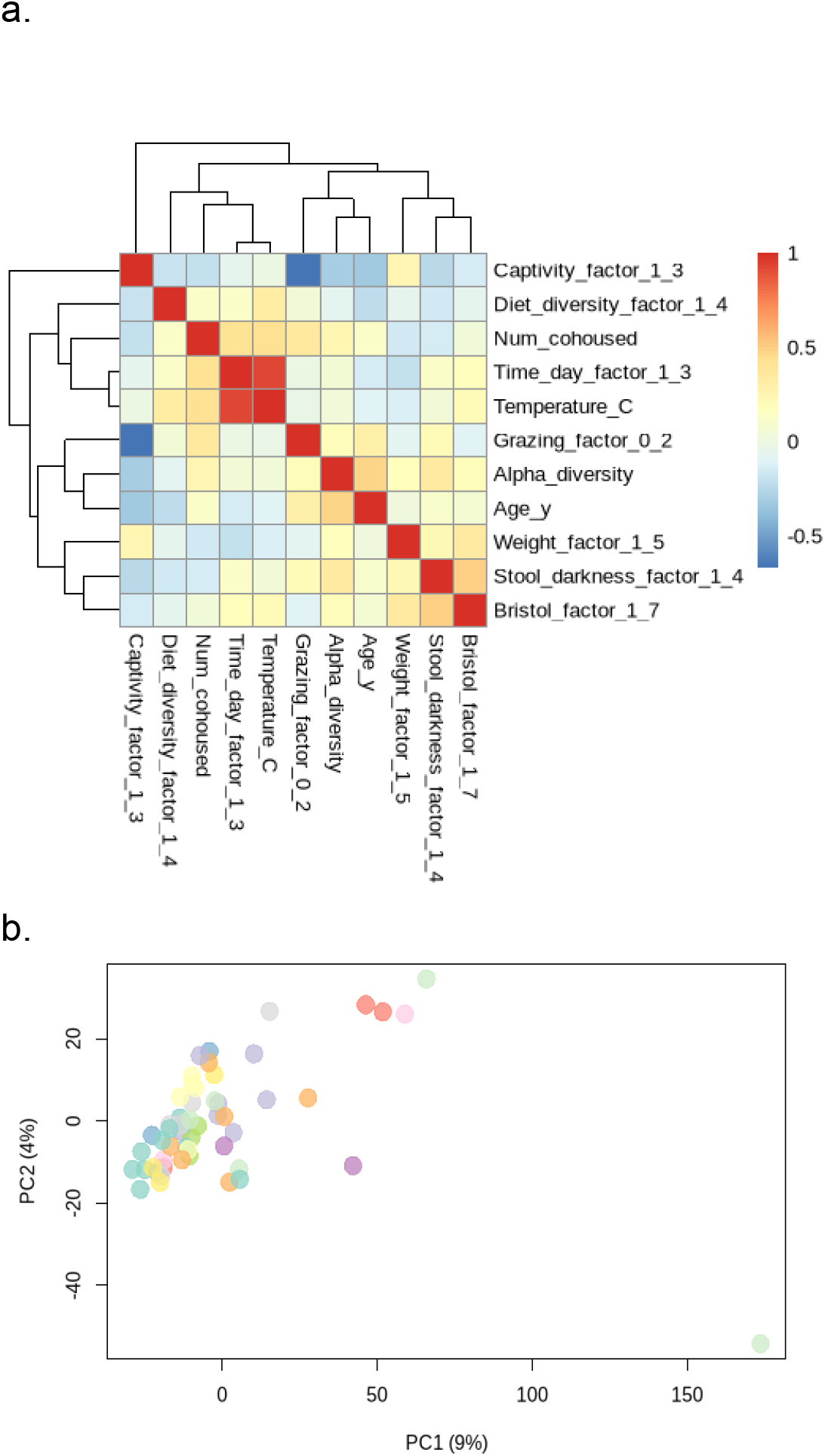
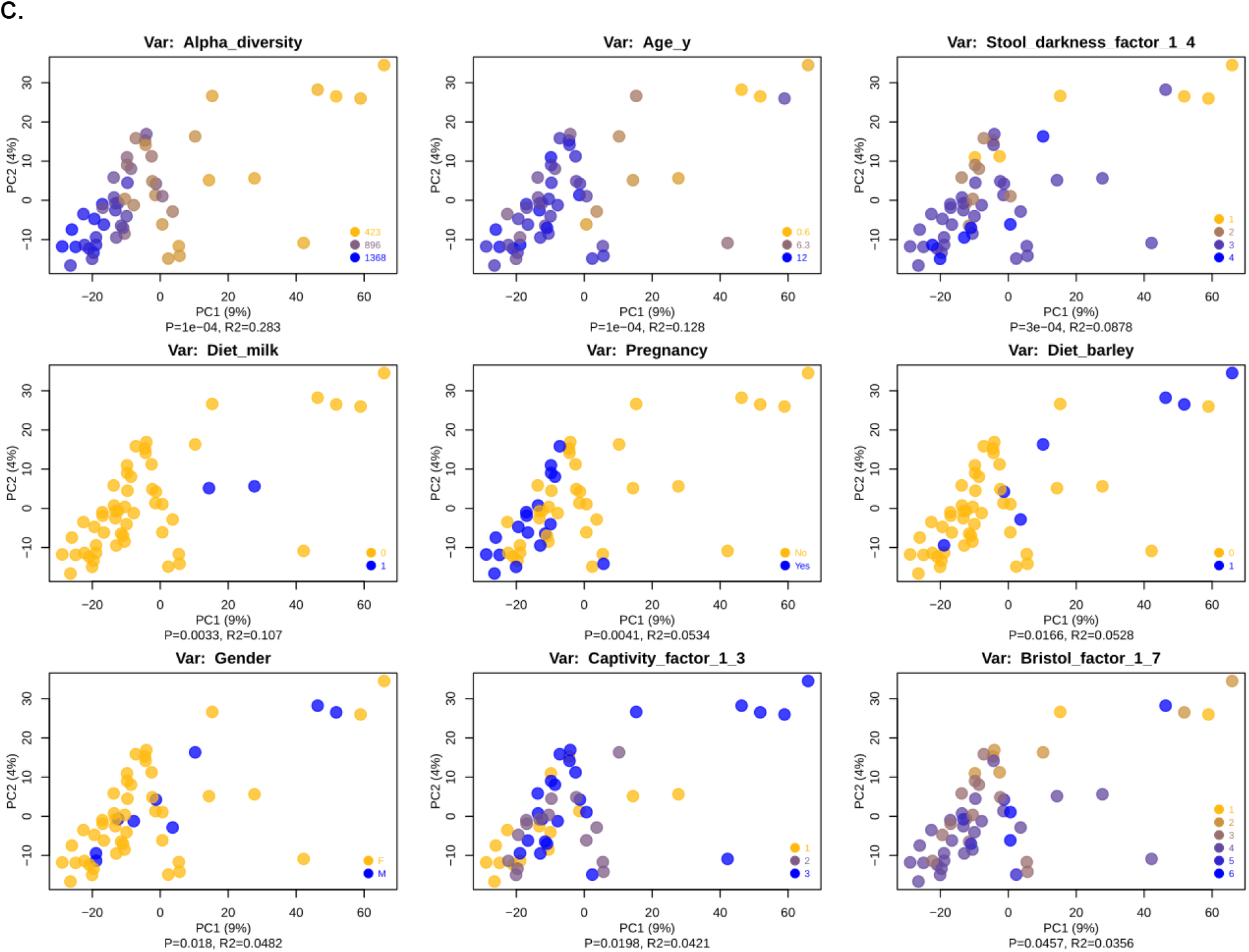
Principles components plots reveal beta diversity distribution colored by various metadata variables (covariates). (a) Symmetric heatmap of pairwise spearman correlations between numerical metadata. (b) shows the entire set of 55 camels colored by herd ID. The outlier in the bottom right is from a camel that is in its third month of life, and was removed (for plotting purposes only) for subsequent beta diversity plots in (c).(c) Beta diversity colored by significantly associated covariates as determined by PERMANOVA p < 0.05. R^2 value (effect size of association) is also displayed under the x axis along with the p-value of the association. For continuous variables, the legend shows the minimum, maximum, and midpoint of the distribution of values plotted. Percentages in axes labels are in terms of percent variance expressed by each axis. The points are in the same place in all plots; only the colors and statistical results differ by metadata variable.

A visual inspection of the herd-labeled beta diversity ordination (Figure 2b) appears to confirm the expectation that camels from the same herd have more similar microbiomes, with a close clustering of like-colored points. A clear outlier microbiome sample is also apparent in the same plot, which was collected from a very young calf (in its third month of life) which was exclusively breastfeeding, unlike any other camel in the dataset (the next youngest is twice its age and consuming solid food).

We tested each metadata variable in the binary and numerical sets using PERMANOVA on the beta diversity distance matrix to determine the significance of the association, and displayed all significant results as colored annotations in Figure 2c. Notably, alpha diversity was a significant driver of beta diversity, forming a clear gradient along PC1, and age followed a similar trajectory (age is also associated with alpha diversity directly; spearman r = 0.61, p = 7.4e-7; pearson rho = 0.48, p = 0.0002). Bristol index and stool darkness followed similar gradients along PC1. Captivity factor shows a reverse trend. Interestingly, although dietary diversity does not produce a significant association, individual dietary components (milk, barley) show some evidence of microbiome clustering. Pregnancy status and gender also produce significant separation. Covariates notably absent from the significantly associative beta diversity results are disease status (we return to this point in later analyses), weight, and number of co-housed camels.

### Differential analysis and machine learning

Relative abundances of genera were used as features to train a random forest model across all metadata variables (Figure 3). Additionally, the top-scoring genera by feature importance were also separately visualized and statistically analyzed using univariate regression and associated statistics. Strikingly, some variables, including all binary dietary features, yielded strong predictive performance (OOB ROC AUC > 0.72), including some features that were not distinguished by earlier analyses of diversity metrics (community-wide). This is especially apparent with the prediction of dietary wheat, which produced a non-significant association with beta diversity (PERMANOVA p=0.13), yet a 0.9 AUC value by random forest prediction using genera relative abundance. Considering wheat in particular is fairly balanced (29 camels are not wheat consumers, and 26 are), it is less likely to be artefactual than some other results with less balanced classes such as milk or grass, where only 3 camels did and didn’t consume these dietary sources, respectively. Milk and grass consumption are also confounded by age and wheat consumption, both of which are highly predictive models on their own.

**Figure 3.**
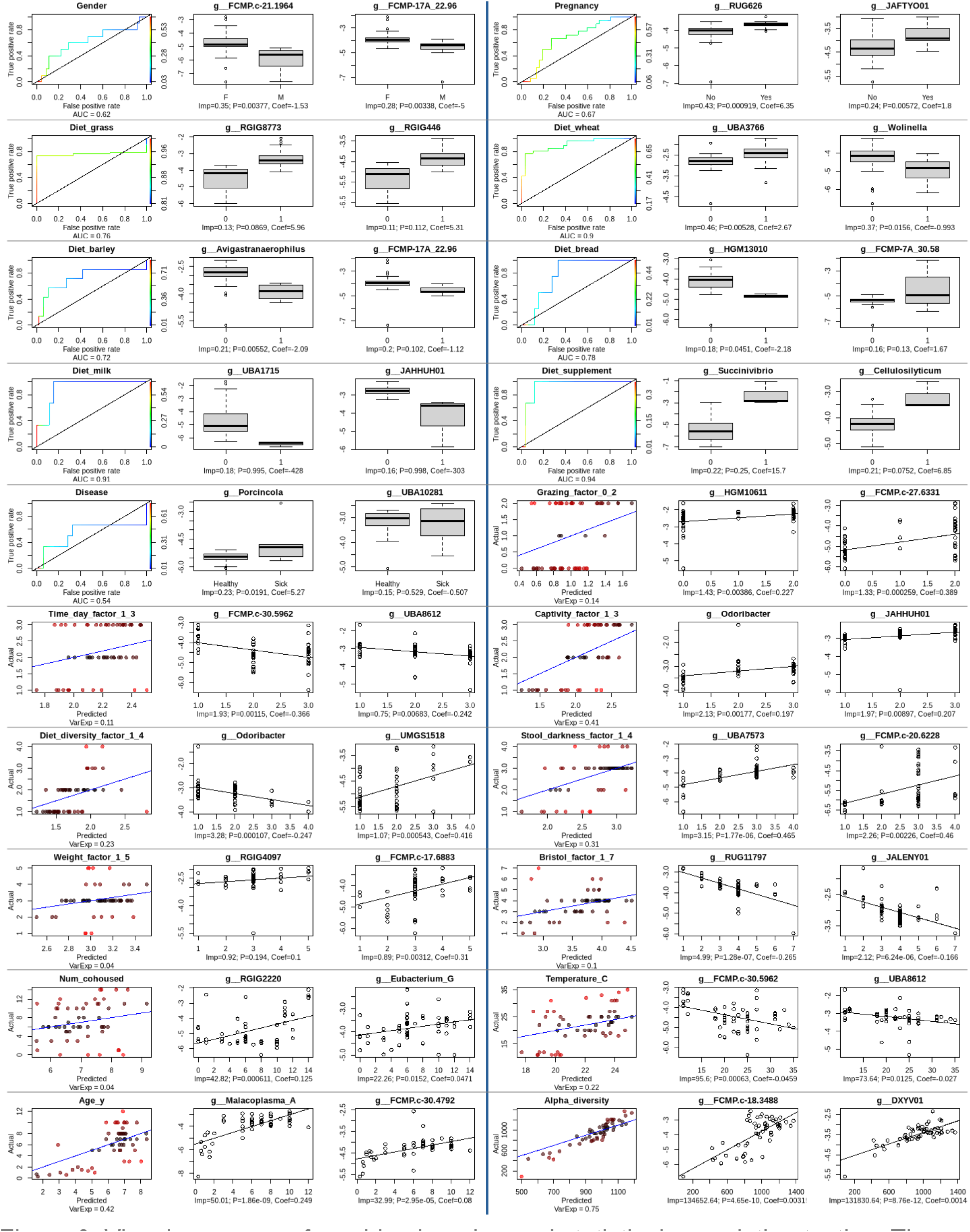
Visual summary of machine learning and statistical association testing. The left-most figure in each panel shows the random forest performance of a model trained on the given variable. Binary variables were used to run random forest models in classification mode, whose performance is conveyed using a colored Receiver Operator Characteristic (ROC) curve, colored by the class probability threshold used for predicting class membership and labeled for the predictive performance based on Area Under the Curve (AUC). The identity (y=x) function has been added, denoting the performance of a hypothetical random model (AUC=0.5). Continuous and ordinal variables were subjected to classification by random forest, which is shown as a scatterplot of the actual value against the out of bag prediction. The identity (y=x) line has been added, denoting hypothetically perfect predictive performance (all predicted values exactly match the true values). Points are colored by their distance from the identify function; more intense red signifies poorer prediction for those values. To the right of each machine learning performance summary plot are two plots showing the log10 relative abundance of the top 2 genera (highest Gini index for the random forest), as either boxplots (classification) or univariate scatterplots (regression) of the genus vs the variable. Each univariate genus plot is labeled with the Gini importance score (“Imp”), as well as both the p-value and coefficient from a univariate model fit (logistic model for binary variables, linear models for the rest).

## Discussion

The Fathi Camel Microbiome Project pilot demonstrates powerful early insights into the diversity and ecological drivers of the Arabian camel microbiome, and lays the groundwork for subsequent analyses and expansion. Although some significant limitations currently exist at this stage, such as small sample size and limited geographical range of sampling, the study design compensates for lack of breadth with greater depth: the sequencing technology (whole-metagenome shotgun) and read depth (110 million total reads per sample) is unprecedented in the literature for the study of camels (and most other livestock), and adopts a genomic analysis approach more common in human metagenome studies. Furthermore, there is considerable potential for the genomic reference databases generated here to more comprehensively characterize future (including shallower) metagenomic studies in camels, reducing the barrier to entry for future work in this agriculturally and culturally important area.

Other limitations include using out-of-bag prediction performance reporting for our random forest models, rather than an explicit cross-validation framework. This is largely due to sample size constraints. Random forests have been previously reported to work well in the high-dimensional low-sample-size domain for microbiome data [19], and out-of-bag prediction performance, which while often representative, may in some cases be optimistic or inaccurate. Nevertheless, the purpose of this approach as implemented in this study is not to create a robust production model, but as a proof-of-concept to highlight promising areas of biological interest that may be worth future follow-ups and secondary validation with a future sampling effort.

Some of the strongest associations uncovered in this analysis relate to age and diet, two interleaving aspects of development that have been shown to be correlated in humans as well [20,21]. The ability of the microbiome to strongly predict dietary components in our study, most notably wheat, is a fascinating glimpse into the ecological ramifications of microbe-food associations writ large. Is it a commensal microorganism that aids digestion of wheat fiber specifically? Or a common plant symbiont that’s merely along for the ride? Such questions would beg longitudinal analysis, interventional studies using dietary swaps, or even a more thorough environmental sequencing of the foodstuff (plant material itself) to address these open questions.

Just as notable, but perhaps not unexpected, is the lack of identifiable association between disease status and microbiome signatures. Although the microbiome has been implicated in numerous diseases in humans and livestock as alluded to in the introduction, we remain underpowered to detect trends in this study due largely to the lack of diseased camels in the populations sampled at time of sampling, as well as more detailed information on the disease etiology. Without finer-grain control as to the nature of the diseases under investigation, our detection of disease signal would be limited to general dysbiosis signatures, which may require significantly larger numbers of matched diseased and healthy camels.

### Future directions

In terms of project scope and expansion, our near-term efforts would seek to expand the sample number to a few hundred camels, as well as add targeted sequencing of camel milk, which is of vital social and economic significance to the region. We are also planning to sample a larger geographical area and explicitly address the question of whether Arabian camel microbiomes have a strong geographic taxonomic signature.

Analytically, we are planning to perform cross-domain (viral, fungal, eukaryotic microbial) identification of microorganisms, as well as attempting to tag consumed plant matter in the fecal DNA. With expanded geographical sampling, we hope to train models to generalize, and assess geographical generalization performance, via a leave-one-region-out approach. Analysis of the results of gene calling, protein clustering, and deep functional annotations of this deep metagenomic data is highly desirable for future work (and indeed, we have already completed much of the computational work underlying this data, as it involves steps which were necessary for other steps in our Methods above; e.g. gene-calling and KEGG ortholog functional annotations are required inputs for CheckM2). Functional annotations may allow us to further explain (or decipher anew) the relationship between microbes and the dietary or environmental features with which they are highly associated in our data; this is also left to future work.

However, due to the high-p low-n dimensionality issues discussed above in the context of the small sample sizes available to us in this initial effort, further increasing the feature space (with genes, proteins, viruses, etc) was deemed risky without a concomitant increase in sample number as planned for the next stage of future work on the FCMP.

## Conclusions

Overall, the current study demonstrates an intensive look into the gut microbiome ecology of the Arabian camel. Utilizing deep metagenomic shotgun sequencing, our study reveals a remarkable amount of novel microbial diversity within 55 sampled camel gut microbiomes. We develop a comprehensive, publicly available database and genomic resource for use in prokaryotic species identification using metagenome-assembled microorganisms. We hope that our microbiome reference database will prove to be a valuable resource for data analysis in camels, as well as in expanding the catalog of global microbial diversity discovered to date.

Our investigation into the correlations between Arabian camel microbiota and various collected covariate metadata exposed some fascinating microbial trends with respect to physical features of these Arabian camels, as well as captured camel gut ecological diversity in broad strokes. Interestingly, our study also provided a tantalizing peek into prospective predictive biomarkers for many of these variables, highlighting the potential to generalize and expand upon these with more samples, and potentially provide clues into the intimate relationship between camels, the microbes they host, the foods they consume, and the environment in which they live.

Furthermore, in using machine learning and confirmatory statistics to highlight the importance of microbial biomarkers (most of which are indeed novel and uncharacterized organisms) for numerous traits, our study highlights the potential to more generally predict camel characteristics, including those we have not yet sampled or recorded, potentially also including those that did not (yet) show predictive capability in our limited sample set. These initial findings, along with future confirmation and deeper understanding thereof, may lead to a more “personalized” perspective in animal lifestyles, including camel husbandry and health care, and potential applications along these lines in other domestic animals as well.

In conclusion, our project has made significant strides in deeply characterizing the species-level microbiome diversity in Arabian camels. It also holds the potential to further advance metagenomic studies in camels and beyond, with a valuable reference database with which to compare results across studies. It uncovered thousands of novel prokaryotic microorganisms without species (and often genus-level) representatives in any existing database, opening up new areas to explore in comparative genomics and systemetology. Most importantly, our study’s early insights and exploratory frameworks will pave the way for future work to reveal the secrets nestled within the bowels of these iconic mammals, including potential future insights into the health and productivity of these camels and beyond.

## Data Availability

All necessary data, including source files, code, runtime parameters, figures are available on GitHub: https://github.com/mubar003/Camel_microbiome_paper Raw metagenomic sequencing data is available from the NCBI sequence read archive (SRA) using BioProject accession PRJNAXXXXX

## Acknowledgment

The authors extend their appreciation to the Deanship of Scientific Research at the University of Tabuk for funding this work through Research No. S-1443-0243. We would like to express our deepest appreciation to Dr. Gabriel Al-Ghalith, whose invaluable expertise and assistance with high-performance computing were instrumental in the successful progression of this project. We extend our sincere gratitude to Abdulrahman Mubaraki for his invaluable assistance during the project period.

